# Design of effective personalised perturbation strategies for enhancing cognitive intervention in Alzheimer’s disease

**DOI:** 10.1101/2023.04.20.537688

**Authors:** Jakub Vohryzek, Joana Cabral, Yonatan Sanz Perl, Murat Demirtas, Carles Falcon, Juan Domingo Gispert, Beatriz Bosch, Mircea Balasa, Morten Kringelbach, Raquel Sanchez-Valle, Giulio Ruffini, Gustavo Deco

## Abstract

One of the potential and promising adjuvant therapies for Alzheimer’s disease is that of non-invasive transcranial neurostimulation to potentiate cognitive training interventions. Conceptually, this is achieved by driving brain dynamics towards an optimal state for an effective facilitation of cognitive training interventions. However, current neurostimulation protocols rely on experimental trial-and-error approaches that result in variability of symptom improvements and suboptimal progress. Here, we leveraged whole-brain computational modelling by assessing the regional susceptibility towards optimal brain dynamics from Alzheimer’s disease. In practice, we followed the three-part concept of Dynamic Sensitivity Analysis by first understanding empirical differences between healthy controls and patients with mild cognitive impairment and mild dementia due to Alzheimer’s Disease; secondly, by building computational models for all individuals in the mild cognitive impairment and mild dementia cohorts; and thirdly, by perturbing brain regions and assessing the impact on the recovery of brain dynamics to the healthy state (here defined in functional terms, summarised by a measure of metastability for the healthy group). By doing so, we show the importance of key regions, along the anterior-posterior medial line, in driving in-silico improvement of mild dementia and mild cognitive impairment groups. Moreover, this subset consists mainly of regions with high structural nodal degree. Overall, this in-silico perturbational approach could inform the design of stimulation strategies for re-establishing healthy brain dynamics, putatively facilitating effective cognitive interventions targeting the cognitive decline in Alzheimer’s disease.

## Introduction

For the delivery of an effective cognitive intervention, the ongoing state of the patient is surely important. For example, it is likely that delivering cognitive intervention to a drowsy person will result in minimal to no improvements. Similarly, neuropathological states such as Alzheimer’s disease (AD) can prevent a more effective cognitive intervention compared to that of a healthy person. The contention here is that neurostimulation can drive the Alzheimer’s brain state to healthier dynamics so that intervention could be more effective.

Alzheimer’s disease is a complex neurodegenerative disease resulting in decline of cognitive functions^1^. Concerted scientific efforts have been developed to study and understand its progression across fields such as genetics, molecular biology, and neuroimaging^2^. These initiatives have generated a lot of valuable data to study, diagnose and analyse the disease progression^1^. Yet, a more mechanistic interpretation is required both for understanding the disease itself^3^ and for predicting the outcomes of potential pharmacological interventions, non-invasive electrical stimulations and cognitive training treatments^4–6^.

Conceptually, we can consider every neurodegenerative or psychiatric condition, such as Alzheimer’s, Schizophrenia and Autism, to be depicted by a condition-specific brain state with altered spatio-temporal dynamics compared to that of a neurotypical (healthy) brain state^5^. This opens avenues towards the diagnosis, prognosis, and potential treatment intervention paradigms^7^. How to best summarise discrete brain states has been the subject of current focus in the field^8^, with the prevailing idea being that a brain state can be described by its spatio-temporal dynamics, understood in terms of functional networks organised in space, waxing and waning in time^9^. This description in turn serves for mechanistic scenarios where in-silico models use information at lower scales (microscopic or mesoscopic) to causally explain the observed changes at the large-scale brain state level^10^. Furthermore, it can serve for stimulation paradigms where mechanistic explanations are relevant in suggesting adequate protocols^4–6^.

In neuroimaging, functional magnetic resonance imaging (fMRI) has revealed statistically significant alterations in brain signals in various conditions, including AD. To this date, multiple studies have demonstrated alterations in AD in terms of resting-state functional connectivity (FC), i.e., how the signals in different brain areas correlate together over time. On one hand, an overall decrease has been detected in functional connections with both hippocampa^l11–13^ and posterior cingulate regions^14,15^. On the other hand, increases in FC were detected between prefrontal cortex and hippocampus^11^, as well as prefrontal and posterior cingulate cortices^14,15^. For these results, a potential compensatory mechanism driven by the prefrontal cortex was suggested, especially in the earlier stages of the disease^16–18^. On the level of intrinsic/resting-state networks, decreases in the Default Mode Network (DMN)^19,20,21^ and increases in the Frontoparietal Network (FPN)^21^ were observed. The DMN results are in line with earlier fMRI task studies^22–25^.

Recently, non-invasive brain stimulation methods targeting the degenerating cortex have been tested to partially reverse the alterations observed in brain activity in AD^26^. However, despite its promising potential, the treatment outcomes of such trial-and-error interventions remain weakly reliable mainly due to a lack of a full mechanistic understanding of brain function and the need for a more principled assessment of the correct perturbations sites to re-establish brain functional connections and potentially drive clinical improvemen^27^. Whole-brain network models, mimicking the dynamic interactions between brain regions at the large-scale, have been used to further understand the mechanisms that drive alterations in functional connectivity^28^. Specifically addressing AD, Demirtas and colleagues used a whole-brain model to show that noisier local dynamics and lateralization, mainly in the left temporal lobe, drive the observed changes in FC^29^. More recently, Perl and colleagues further demonstrated increased stability in the simulated activity of hippocampal and insular regions when brain atrophy was considered in the whole-brain model, further suggesting perturbation protocols to drive in-silico recovery towards the healthy state^30^.

Here, our goal was to quantify the level of regional susceptibility to rebalance brain dynamics from the different AD stages to the optimal brain state. To do so, we followed the three-part concept of Dynamic Sensitivity Analysis^5^, where we first summarised the large-scale functional organisation of Healthy Controls (HC), Mild Cognitive Impairment due to Alzheimer’s Disease (MCI) and mild dementia due to Alzheimer’s Disease (AD) cohorts in terms of global brain connectivity (GBC), metastability and synchrony. Secondly, we built computational models for all individuals in the MCI and AD conditions. Lastly, we stimulated the brain regions and assessed the impact on the recovery of brain dynamics to the healthy state (here defined by the measure of metastability of the HC group). We derived Perturbation Effectivity for Recovery (PER) as the difference between the simulated and target metastability. Our hypothesis was that both MCI and AD conditions will have similar in-silico regional perturbation profiles but the intensity of perturbation needing to drive the transition to the healthy state will be less in the MCI group.

## Material and methods

### fMRI data

We used a fMRI dataset from a cohort of 97 participants from the Hospital Clinic de Barcelona, Barcelona, Spain, classified into 3 groups: 58 HC subjects aged 60.72±6.99 years old (mean±standard deviation, SD), 20 women; 23 MCI patients aged 69.73±7.77 y.o., 9 women; and 16 AD patients aged 65.00±9.98 y.o., 7 women. The fMRI signals were recorded over 10 minutes with a sampling rate of 2 seconds and averaged within N=90 regions of interest representing cortical and subcortical brain regions defined using the AAL atlas^31^. A more detailed description of the acquisition and pre-processing steps can be found in^29^. Clinical and neuropsychological assessments were recorded to quantify the APOE4 carrier status, the Aβ1–42, p-tau and t-tau values in CSF, and memory test scores in terms of Buschke AL, AT, RDL and RDT.

After the pre-processing of the fMRI timeseries, we first demeaned the regional timeseries and band-pass filtered the data in a narrow range from 0.008 to 0.08 Hz, to exclude physiological noise confounds, known to be less relevant in this range. The phase of the fMRI signals in this narrow-band was subsequently estimated using the Hilbert transform^32^. The Hilbert transform expresses the regional timeseries in terms of a time-varying phase *θ*(*t*) and amplitude *A*(*t*) as follows *x*(*t*) = *A*(*t*) \**e*^*i*θ(*t*)^. The three first and last timepoints of the regional timeseries were excluded due to the boundary artefacts introduced by the Hilbert transform.

### fMRI measures

We considered in this work measures of fMRI signals to describe brain states that were previously shown to be significantly altered in patients with Alzheimer’s disease^29^, namely Global Brain Connectivity (GBC), metastability and synchrony (**Figure 1A**). These measures were calculated for each fMRI scan and compared statistically between groups. The measure of metastability was subsequently used to adjust the parameters of the whole-brain network model to approximate the brain dynamics of MCI and AD participants (**Figure 1B**) and evaluate the optimal stimulation sites to approximate HC brain state.

**Figure 1.**
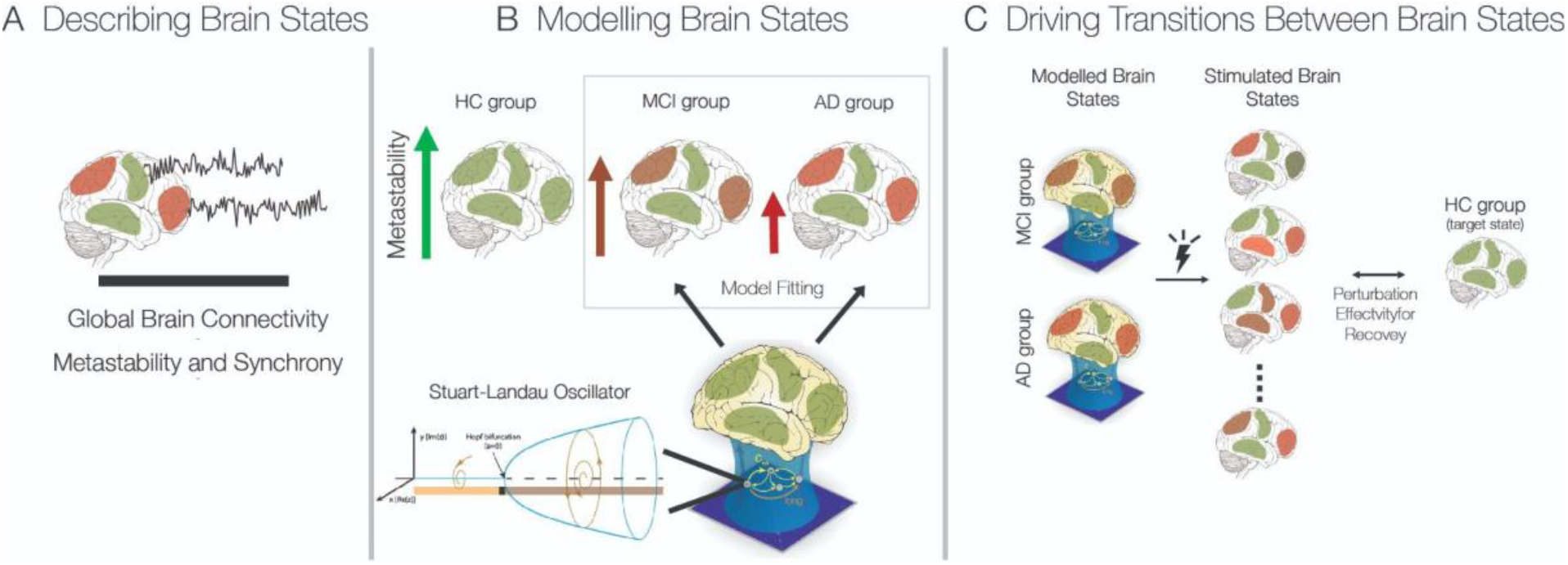
Study Overview. A) Describing Brain States. Each brain state – here describing healthy, MCI and AD brain dynamics can be summarised with various measures. We measured Global Brain Connectivity, Metastability and Synchrony. **B) Modelling Brain States**. We built individualised models for the MCI and AD group by minimising the difference between the empirical and model Metastability measure. **C) Driving Transition Between Brain States i.e. “Dynamic Sensitivity Analysis”**. We performed in-silico bilateral perturbations to achieve the optimal protocol for a transition between the diseased brain states and the healthy (target) brain state (described by the mean empirical metastability for the healthy group).

### Global Brain Connectivity

For the computation of global brain connectivity (GBC), we first estimated the NxN functional connectivity matrix as the Pearson correlation between the *N=90* band-pass filtered fMRI timeseries. Then, global brain connectivity was defined as the structural nodal strength

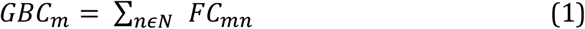

here *FC*_*mn*_is correlation between regions ***m*** and ***n***, and *N* is the number of regions. Furthermore, we define the mean GBC, ⟨*GBC*⟩, as the average across brain regions.

### Synchrony and metastability

The measure of synchrony describes the average phase coherence of fMRI signals across an entire scan, and metastability quantifies how much the levels of phase coherence fluctuate in time. To evaluate the phase coherence over time, the Kuramoto Order Parameter is calculated at each instant of time as follows:

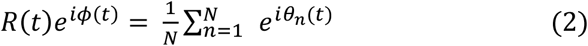

where θ_*n*_(*t*) is the instantaneous phase of each region **n** at time **t**. *R*(*t*) is the order parameter amplitude quantifying the overall coherence of all the regional phases at time **t**, such that R = 0 when the system is fully desynchronized, and R = 1 when it is fully synchronised. Synchrony and metastability are defined as the mean and standard deviation of the Kuramoto Order Parameter over time, respectively^33^. Intuitively, metastability is low when the level of synchrony of the system is stable in time (either high or low synchrony), with maximal metastability obtained when the system fluctuates between periods of low and high synchrony over time.

### Statistical comparison between conditions

The statistical analysis for the group difference at the empirical level was quantified with unpaired two-sided t-test statistics (significance threshold of 0.05). We report corrected (Bonferroni test) and uncorrected p-values visualised by a green and red star respectively. The statistical testing for non-stimulated and optimal PER values were performed with the same analysis for two-sided paired t-test statistics. The correlation analysis between PER values and physiological and cognitive biomarkers was performed with Pearson’s correlation and the results are reported for uncorrected p-values. The relationship of optimal PER values across brain regions and their structural hubness was also quantified with Pearson’s correlation.

## Whole-brain Computational Model

### Structural Connectivity

For the whole-brain model, we used a group-based structural connectivity matrix derived for the same *N=*90 brain areas. The group-based structural connectome was obtained from previously scanned 16 healthy young adults recruited online at Aarhus university (5 females, age (mean±SD): 24.7 ± 2.54 y.o.). The details of the dataset and pre-processing steps for the derivation of the structural connectome from diffusion tensor imaging (DTI) are described in^34^. Individual undirected structural connectivity matrices *Cnp* were defined, where *C* encodes the connectivity weights in proportion to the number of sampled fibers between regions **n** and **p**. To obtain the group structural connectome, the individual connectomes were averaged across the sixteen subjects.

### Model equations

We used a whole-brain network model to approximate the brain dynamics of MCI and AD participants, consisting in a system of coupled oscillators, each oscillating in the ultra-slow frequency range of fMRI signals, and coupled together according to the weights derived from the structural connectivity matrix^35,36^. Such whole-brain models have been previously shown to reflect the emerging brain dynamics in fMRI^35–38^.

The oscillatory dynamics of each brain region (i.e., each node n in the network) is modelled using the Stuart-Landau equation, which is a canonical model to describe the behaviour of an oscillator with a supercritical Hopf-bifurcation, i.e., depending on the parameters, it can exhibit either damped or self-sustained oscillations^39^.

In Cartesian coordinates, the oscillatory dynamics of each uncoupled region of interest (n) is described as follows

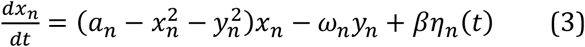

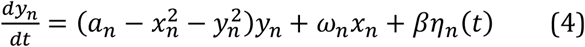

where *η*_*n*_(*t*) represents added gaussian noise with standard deviation of *β* = 0.02, *a* is the bifurcation parameter that positions each brain region at different dynamic regimes (for *a* = 0 at the bifurcation point, *a* < 0 in the fixed point dominated by *βη*_*n*_gaussian noise, and *a* > 0 in the stable limit cycle with oscillations at the intrinsic frequency of *f*_*n*_= *ω*_*n*_/2π Hertz, c). The values of *ω*_*n*_are derived for each region of interest n and for each patient from the empirical fMRI data by taking the peak frequency of the gaussian-smoothed power-spectrum of the band-pass filtered (0.04-0.08Hertz) fMRI signals. Then, we coupled the regional equations according to the structural connectivity (C) that approximates the large-scale white-matter connectivity of the human cortex^35,40^. The coupling term, connecting the different regional equations in the whole-brain network, is modelled as a common difference coupling (the linear term of a general coupling function) and weighed by the matrix C. It is to be noted that we don’t consider the other non-linear terms of the Taylor expansion of the coupling term^41,42^. Then, the equations 2 and 3 become:

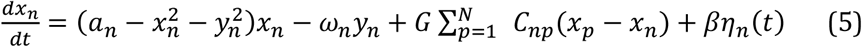

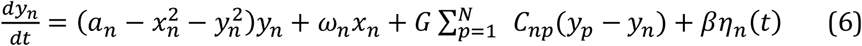

with *C*_*np*_(*x*_*p*_ − *x*_*n*_) describing the difference coupling between region ***p*** and ***n*** weighted by its structural weight *C*_*np*_. The matrix C was rescaled for parameter range consistency with previous works as follows < *C* > = 0.2^35,36^. The variable ***G***, representing the global coupling strength, is the free parameter used for fitting the whole-brain models to the approximate metastable dynamics of fMRI signals in each patient. The bifurcation parameter ***a*** was set to -0.02 in the simulations. The dynamic variable *x*_*n*_represents the simulated fMRI signal for every node ***n***^35,36^.

### Objective Function

For the model fitting we calculated the metastability of the simulated fMRI signals obtained for a range of the free parameter ***G*** (from G = 0.01 to G = 0.5). To compare the measures of the simulated signals with the empirical ones, we calculated the absolute difference between the measures of synchrony, metastability and mean GBC. To compare the GBC of each brain region, we calculated the mean squared error between simulations and empirical values. We fitted the model for each measure separately to show their convergence toward an optimal working point for each patient in every group. Then, for each patient we used the global coupling G associated to the minimum of the absolute difference of metastability.

### Driving transitions between bran states

For the perturbation protocol, we used the bifurcation parameter ⍰_⍰_ to alter bilaterally regional dynamics either towards a more noise-driven (⍰_⍰_<0) or oscillatory regime (⍰_⍰_>0). After each stimulation to bilateral areas, we calculated the absolute difference between the metastability of the simulated signals and the average metastability measured empirically in the HC group. For both the AD and MCI groups, we systematically perturbed the bilateral regions (in total 45 pairs of regions) and compared the metastability of the simulated signals under perturbation to that of the target state (**Figure 1C**).

## Results

### Describing Brain States across conditions

We first assessed differences in fMRI signals between HC, MCI, and AD conditions to appropriately characterise the altered dynamics in the different brain states so those metrics can be recovered in the models. Mean Global Brain Connectivity was altered between HC and AD group as well as HC and MCI group and non-significant between MCI and AD groups (unpaired uncorrected t-test, mean GBC: HC vs. AD p-val = 0.0045, HC vs. MCI p-val = 0.0169., MCI vs AD p-val >0.05, **Figure 2A top left**). This can be further appreciated in **Figure 2A top right**, where the GBC values for individual regions are rank ordered, clearly indicating a decrease in both MCI and AD stages for regions with high values of GBC in the HC group. We further calculated Metastability (unpaired uncorrected t-test, HC vs. AD p-val = 0.0045, HC vs. MCI p-val = 0.0099., MCI vs AD p-val = n.s., green star <0.05 Bonferroni corrected, **Figure 2B**), and Synchrony (HC vs. AD p-val = 0.0040, HC vs. MCI p-val = 0.0096, MCI vs AD p-val = n.s., green star <0.05 Bonferroni corrected, **Figure 2C**) as measures of the collective dynamics of brain regions.

**Figure 2.**
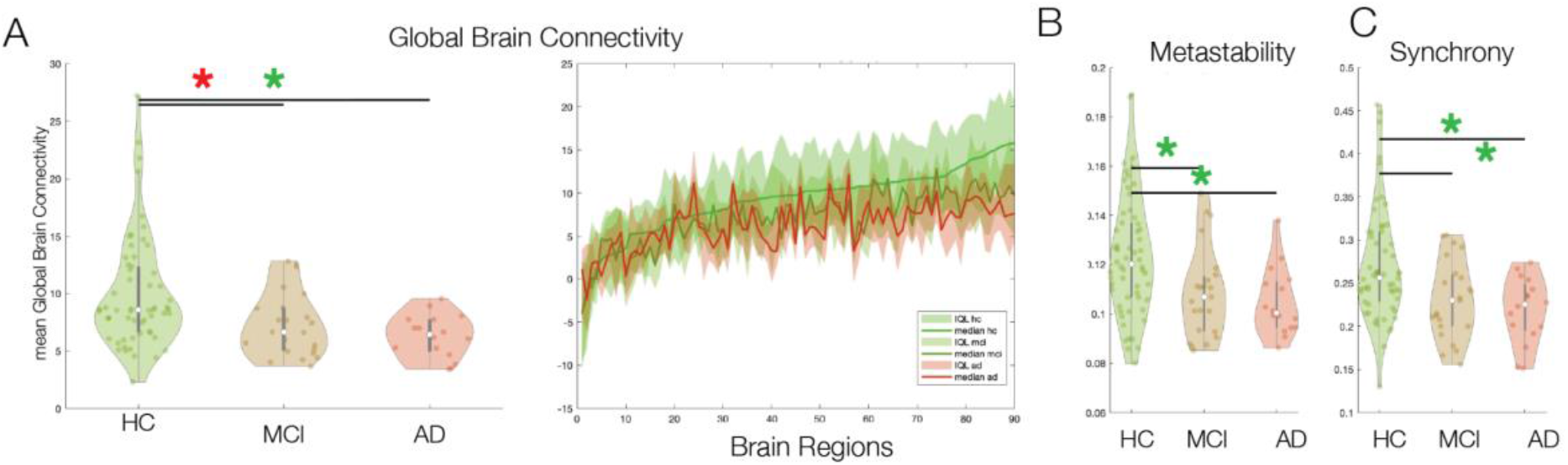
Characterisation of Brain States. **A) Left** Mean Global Brain Connectivity (unpaired uncorrected t-test, HC vs. AD p-val = 0.0045, HC vs. MCI p-val = 0.0169, MCI vs AD p-val = n.s., green star <0.05 Bonferroni corrected, red star <0.05 uncorrected). **Right** Ranked Global Brain Connectivity showing collapse of the functional hub regions in MCI and AD groups. **B)** Metastability (unpaired uncorrected t-test, HC vs. AD p-val = 0.0045, HC vs. MCI p-val = 0.0099., MCI vs AD p-val = n.s., green star <0.05 Bonferroni corrected). **C)** Synchrony (HC vs. AD p-val = 0.0040, HC vs. MCI p-val = 0.0096, MCI vs AD p-val = n.s., green star <0.05 Bonferroni corrected).

### Modelling Brain States for Alzheimer’s

For the model fitting, we chose metastability as a representative measure of brain spatio-temporal dynamics due to its strong ability to separate the AD and MCI from the HC group and due to this measure’s reliability in Alzheimer’s Disease^29^. As such, we fitted each patient’s whole brain model by minimising the absolute difference between empirical and simulated metastability for the free parameter of global coupling G (**Figure 3A**). The optimal fit was different for each group, with an average global coupling G = 0.200 ± 0.076 for the HC group, G = 0.178 ± 0.074 for the MCI group and G = 0.134 ± 0.043 for the AD group. The simulated metastability values for each subjects at optimal G were ⍰ ⍰ ⍰ ⍰ ⍰ ⍰ ⍰ ⍰ ⍰ ⍰ ⍰ ⍰ ⍰ _⍰ ⍰ ⍰_= 0.125 ± 0.024 for the HC group, ⍰ ⍰ ⍰ ⍰ ⍰ ⍰ ⍰ ⍰ ⍰ ⍰ ⍰ ⍰⍰_⍰ ⍰ ⍰_ = 0.114 ± 0.022 for the MCI group and ⍰ ⍰ ⍰ ⍰ ⍰ ⍰ ⍰ ⍰ ⍰ ⍰ ⍰ ⍰⍰_⍰ ⍰ ⍰_ = 0.102 ± 0.018 for the AD group (**Figure 3B-D**).

**Figure 3.**
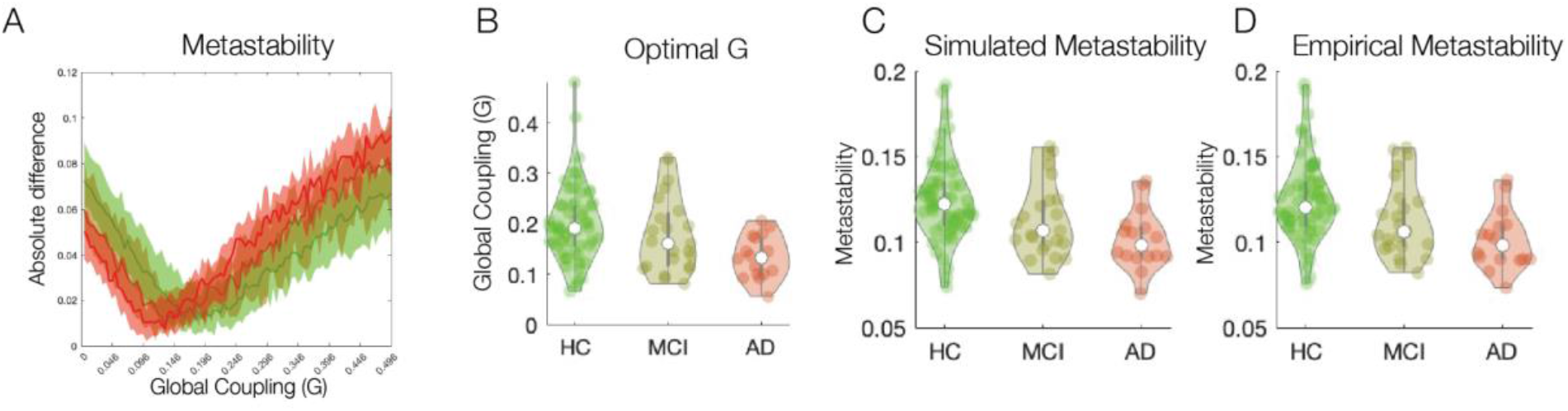
Model Fitting. **A)** Fitted personalised whole-brain models for HC, MCI and AD groups to the measure of Metastability. **B)** The Global coupling parameter G for each group (G = 0.200 ± 0.076 for the HC group, G = 0.178 ± 0.074 for the MCI group and G = 0.134 ± 0.043 for the AD group). **C)** The simulated metastability values for each group at optimal G were metastability_sim_ = 0.125 ± 0.024 for the HC group, metastability_sim_ = 0.114 ± 0.022 for the MCI group and metastability_sim_ = 0.102 ± 0.018 for the AD group. **D)** Empirical metastability reported here for visual comparison to the simulated values of metastability of each group.

### Driving Transitions between Alzheimer’s and a healthy brain state

To assess how the MCI and AD groups can be driven in-silico to a more optimal healthy brain dynamics, we used Dynamic Sensitivity Analysis to rebalance the MCI- and AD-fitted models to the healthy target state as defined by the mean metastability of the HC group. For the AD group, we systematically perturbed the bilateral regions (45 pairs of regions) and compared the metastability of the simulated signals under perturbation to that of the target state (**Figure 4A** left). Perturbation with a negative bifurcation parameter didn’t improve the difference of metastability between target state and simulated AD state under stimulation, whereas perturbation with a positive bifurcation parameter in specific pairs of bilateral brain regions resulted in a minimisation of the difference, indicating a recovery of the metastable dynamics. We highlight regions in green to be driving the improvement of the AD model at an optimal stimulation intensity of a = 0.24 (**Figure 4A right**).

**Figure 4.**
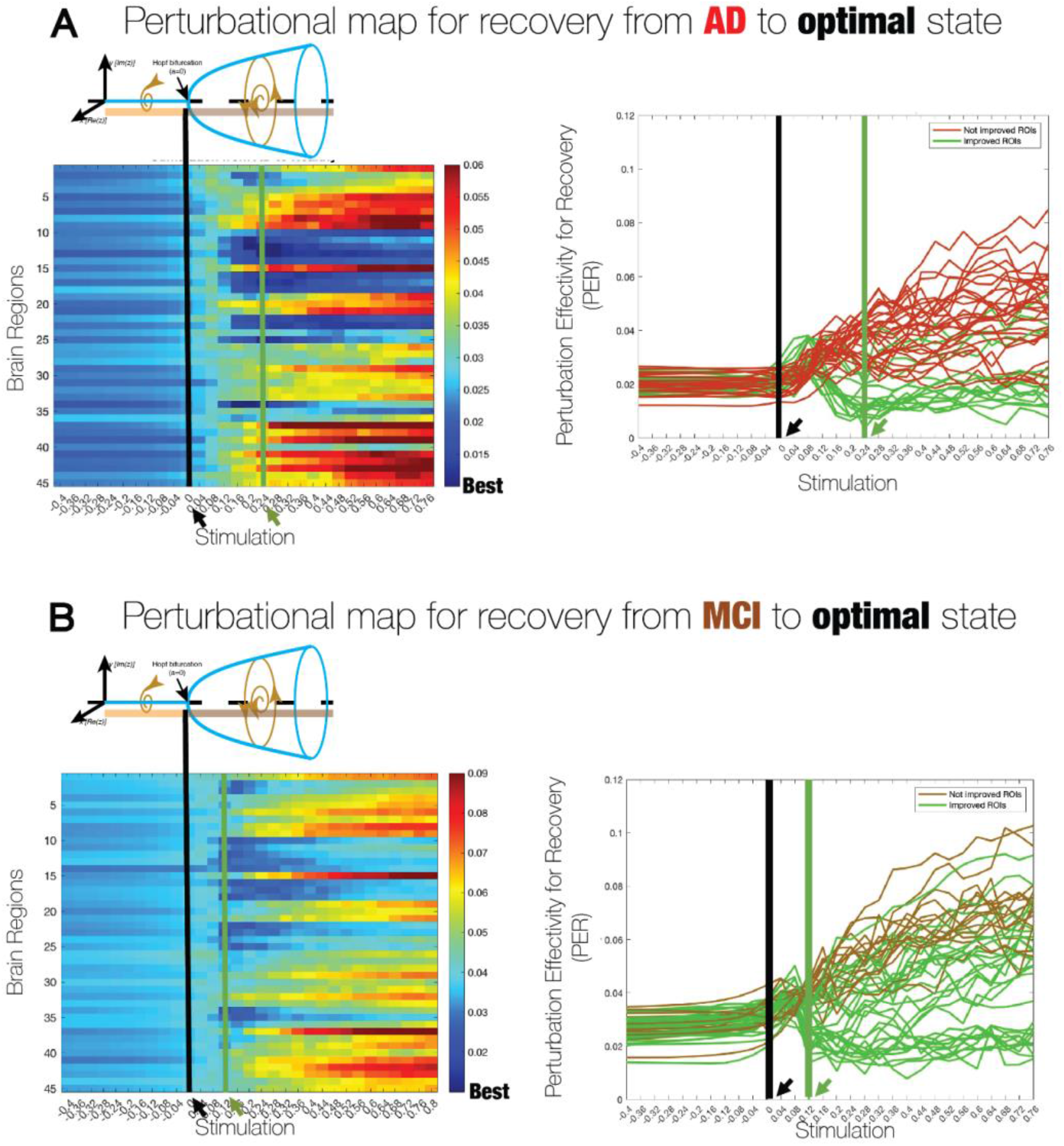
Results of the Stimulation Protocol: **A - Left**: Perturbation brain map for transition between AD to HC for varying stimulation intensity. Optimal stimulation, defined as the mean minimum across brain regions, was achieved at positive (synchronous) stimulation of 0.24 (green line). Black line signifies the fitted model to the AD group before stimulation. **A - Right**: Perturbation Effectivity for Recovery (PER) shows a subset of regions (green) minimising the distance to the optimal metastable dynamics. Again the green line signifies the optimal perturbation across all brain regions and black line indicates the model fit before stimulation. **B - Left**: Perturbation brain map for transition between MIC to HC for varying stimulation intensity. Optimal stimulation, defined as the mean minimum across brain regions, was achieved at positive (synchronous) stimulation of 0.12 (green line). Black line signifies the fitted model to the MCI group before stimulation. **B - Right**: PER shows a subset of regions (green) minimising the distance to the optimal metastable dynamics. Again the green line signifies the optimal perturbation across all brain regions and black line indicates the model fit before stimulation.

When we plotted these regions on a brain surface, a pattern of regions along the anterior-posterior midline axis was observed with the precuneus, the cuneus, the anterior, middle and posterior cingulate cortices, the frontal superior medial gyrus, the frontal medial-orbital gyrus, the calcarine and the olfactory gyrus being among the most statistically significant (**Figure 5A-B top**, red: pval<0.005, orange: pval<0.01, yellow: pval<0.05, white: pval<n.s.). We introduced the same analysis to the MCI-modelled brain state. As in the AD case, perturbation with negative bifurcation parameters didn’t improve the difference in metastability to the target state, whereas bilateral perturbation with a positive bifurcation parameter in specific brain regions resulted in minimisation of the difference (**Figure 5A-B bottom**). Notably, the optimal stimulation intensity to restore metastability in-silico was lower for the MCI group (a = 0.12) compared to the AD group (a =0.24). This is in line with the hypothesis that the MCI brain state requires less stimulation for recovery towards the target state. Again, we highlight regions in green which drive the improvement of the MCI model at an optimal stimulation intensity of a = 0.12 (**Figure 5C top**). In this instance, the frontal superior medial gyrus, the precuneus and the cuneus are the most statistically different brain regions. In addition, as in the AD case, a pattern is observed with all the anterior and posterior cingulate cortices, calcarine and rectus gyrus, as well as the frontal medial-orbital gyrus and the superior motor area (**Figure 5C bottom**, red: pval<0.005, orange: pval<0.01, yellow: pval<0.05, white: pval<n.s.).

**Figure 5.**
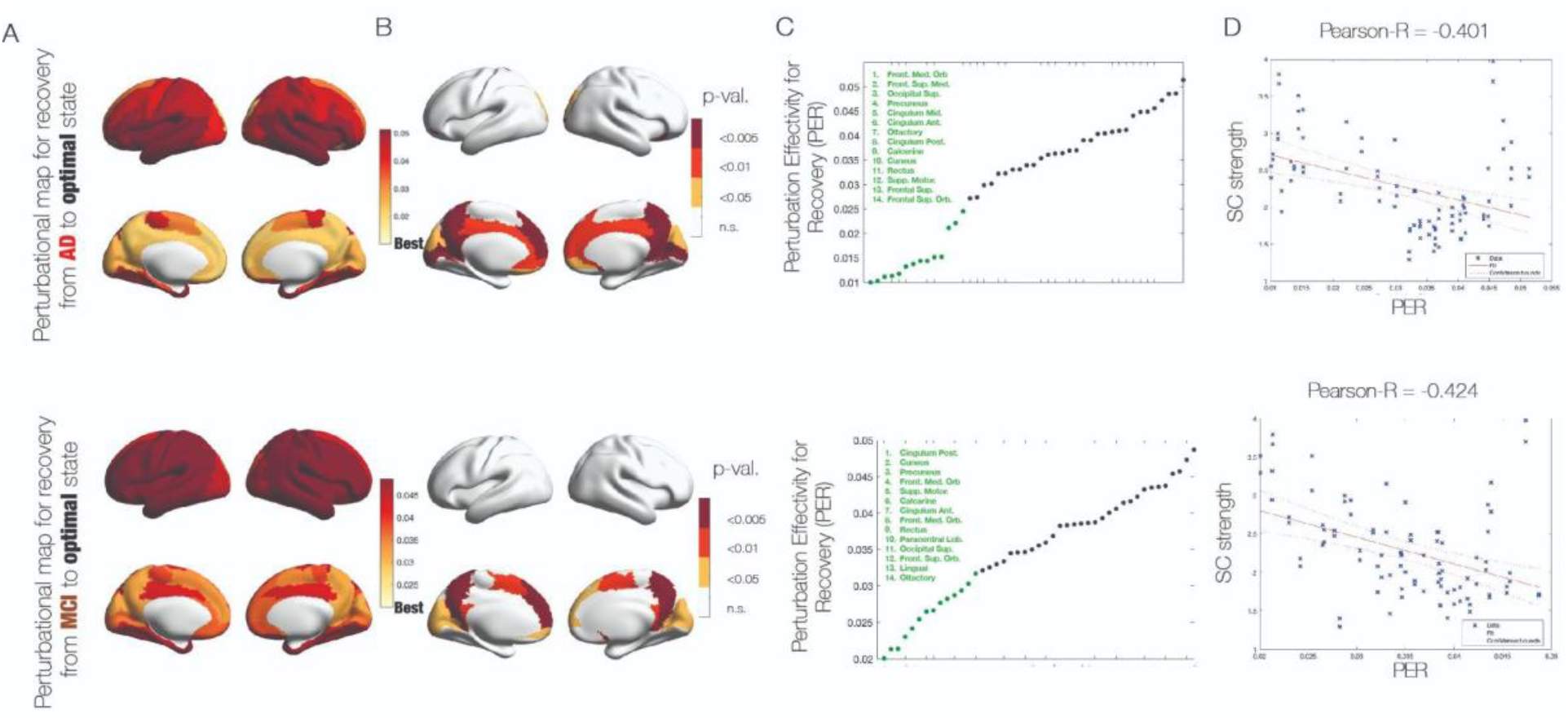
Results of the Stimulation Protocol: **A)** Top: Stimulation brain map for transition between AD to HC at positive (synchronous) stimulation of 0.24. Bottom: Stimulation brain map for transition between MCI to HC at positive (synchronous) stimulation of 0.12 **B)** Top: Paired t-test statistics between the AD (original) and target state (HC). Bottom: Paired t-test statistics between the MCI (original) and target state (HC). Colours represent different statistical thresholds - burgundy p-val<0.005, orange p-val<0.01, yellow p-val<0.05 and white p-val = n.s. **C)** Top: Rank ordered region of AD group by the distance to the HC (target) state. Bottom: Rank ordered region of MCI group by the distance to the HC (target). The green dots represent the strongest 14 regions enabling the transition to the healthy state. **D)** Top: Correlation between the nodal distance in AD group to the HC (target) state and structural nodal strength derived from the structural connectome. Bottom: Correlation between the nodal distance in MCI group to the HC (target) state and structural nodal strength derived from the structural connectome.

### Correlation with structural nodal degree

The propensity of a brain region to rebalance metastable dynamics correlates with structural nodal degree, i.e., how strongly individual regions are connected within the structural network, suggesting that targeting brain hubs has a higher chance of driving the optimal outcome (AD to HC corr = -0.401 p-val = 0.0001, MCI to HC corr = -0.4235 p-val <0.00003). While it might be intuitive that regions with higher connectivity will be more impactful to the overall state of the in-silico brain dynamics, this effect seems to reflect this phenomenon only partially as there are also regions with high structural nodal degree but low influence on the improvement of the aberrant brain dynamics (**Figure 5D**).

### Relation with Physiological and Cognitive scores

To investigate how well the PER measure relates to the physiological and cognitive scores of individual AD patients, we analysed the relationship between the PER measure and the APOE4 allele carrier, CSF and cognitive biomarkers. We computed the correlation between the PER values at the optimal stimulation for the AD group, i.e., how well did the patient’s brain dynamics approximate the optimal healthy brain dynamics, and the APOE4 carrier status and CSF biomarkers. We chose the Superior Frontal Gyrus, Precuneus and Posterior Cingulate Cortex for the exploratory analyses (the statistical significance between the non-stimulated PER (stim=0) and the optimal PER at stim of 0.24 for the three regions was p-val = 0.006, 0.0056, 0.009, respectively). We found a significant correlation of the Superior Frontal Gyrus with the t-tau (pearson’s r = -0.51, p = 0.044) and p-tau (pearson’s r = -0.59, p = 0.016) concentration, the CSF biomarker index (pearson’s r = -0.50, p = 0.049) and Buschke AL cognitive score (pearson’s r = - 0.52, p = 0.040) (**Figure 6A**). Posterior Cingulate Cortex values correlated also with the Buschke AL cognitive score (pearson’s r = -0.50, p = 0.048, **Figure 6B**), while precuneus had all correlation comparisons non-significant (**Figure 6C**). Interestingly, this demonstrates that the amount of in-silico recovery alone cannot determine physiological and cognitive biomarkers, but that the specific anatomic site is important.

**Figure 6.**
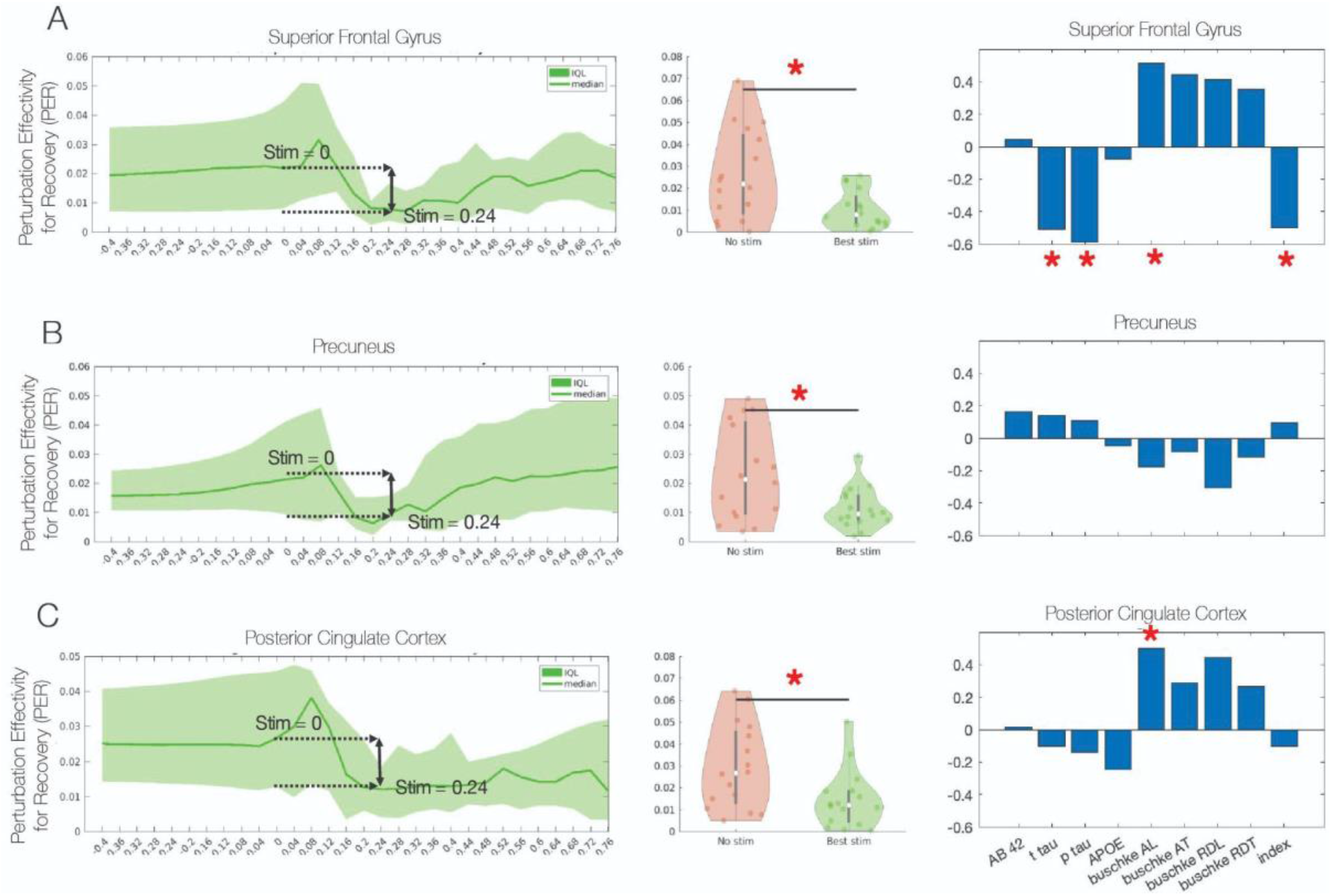
Regional perturbation results and physiological and cognitive scores: **A)** Left: Perturbation map for Frontal Superior Medial Gyrus. Stim = 0 signifies the unperturbed metastability distance of the individual patients in the AD group compared to the average metastability of HC group. Middle: Statistical significance between the non-stimulated (stim=0) perturbational distance at stim = 0.24 (p-val = 0.006, paired t-test). Right: Correlations between patient’s cognitive scores and perturbational distance at the optimal stimulation (stim = 0.24). Significant Pearson’s correlation between t-tau, p-tau, buschke AL and index (Pearson’s R = -0.51, -0.59, 0.52, -0.50 and p-val = 0.044, 0.016, 0.040, 0.049 respectively). **B)** Left: Perturbation map for Precuneus. Stim = 0 signifies the unperturbed metastability distance of the individual patients in the AD group compared to the average metastability of HC group. Middle: Statistical significance between the non-stimulated (stim=0) perturbational distance at stim = 0.24 (p-val = 0.0056, paired t-test). Right: There was no significant correlation for Precuneus. **C)** Left: Perturbation map for Posterior Cingulate Cortex. Stim = 0 signifies the unperturbed metastability distance of the individual patients in the AD group compared to the average metastability of HC group. Middle: Statistical significance between the non-stimulated (stim=0) perturbational distance at stim = 0.24 (p-val = 0.009, paired t-test). Right: Correlations between patient’s cognitive scores and perturbational distance at the optimal stimulation (stim = 0.24). Significant Pearson’s correlation between CSF index (Pearson’s R = -0.50 and p-val = 0.048 respectively).

## Discussion

In this work, we used whole-brain models to design non-invasive stimulation strategies for re-establishing healthy brain dynamics, and facilitation of effective cognitive interventions in Alzheimer’s disease. Following the concept of Dynamic Sensitivity Analysis, we quantified the empirical differences between the HC, MCI and AD groups in terms of their metastability alterations, and fitted subject-specific whole-brain models to the altered brain states of MCI and AD^5^. Using an in-silico stimulation protocol and the Perturbation Effectivity for Recovery index, we were able to rank brain regions according to their proclivity to drive an increase in metastability levels from the levels detected in MCI and AD brain states towards the average metastability detected in healthy controls. These were mainly along the brain’s medial axis both in the anterior and posterior parts. The required intensity of stimulation for successful outcome was lower for the MCI brain state suggesting less impacted brain dynamics compared to the AD brain state. Furthermore, the proclivity of a region to rebalance the dynamics to healthier brain state was correlated with the structural nodal degree.

Using in-silico perturbations as a dynamic sensitivity analysis, one can understand brain state differences from a dynamic system perspective. Such approach complements traditional functional connectivity analysis, where brain states are differentiated via signal detection theory. This is achieved by considering a stimulation scenario where a regional proclivity to rebalance brain dynamics to a target state is quantified. Here, we showed that regions along the medial axis, belonging mainly to the anterior and posterior default mode network, have the greatest impact on rebalancing brain dynamics towards healthier brain state. In fact, this is in line with previous literature demonstrating regions of the default mode network to be impacted in AD stages^19,20,21^. Interestingly, medial regions of the fronto-parietal network are also implicated in the possible rebalancing towards the optimal target state. Indeed, previous literature has shown increases in the activation of FPN in AD^21^. The fact that dynamic sensitivity analysis is agnostic towards the direction of the activity alterations as demonstrated here for DMN and FPN, might reflect the fact that as long as regional alterations exist, they will be reflected in the ability of the system to rebalance back towards a healthier target state.

Beyond dynamic sensitivity analysis, such in-silico perturbation protocols can be used for non-invasive brain stimulation with tools such as Transcranial Magnetic Stimulation (TMS) and transcranial Direct and Alternating Current Stimulation (tDCS and tACS)^43^. Empirical studies have shown the feasibility of non-invasive stimulation in AD for effective cognitive intervention. Thus far, most studies have focused on the stimulation of frontal and parietal lateral regions^6,44,45^. Indeed, this is in line with the results here showing that regions with highest impact to rebalance healthy brain dynamics lie in the frontal and posterior regions. Although, our analysis indicated that regions in frontal and posterior areas along the medial line are the ones more likely to drive the change. Taken together, such empirical and simulated synthesis might be achieved through left-right lateral stimulation where both lateral and medial frontal and parietal regions are impacted as opposed to anterior-posterior stimulation.

Here, we focused on the intrinsic stimulation paradigm where the dynamical regime of the regional Hopf model is altered either towards oscillatory or noise-driven regime. In this respect such stimulations can be thought of as phenomenological. Alternatively, such in-silico modelling paradigms can describe an external perturbation protocol whereby parameters of driving oscillations such as the frequency and amplitude are used to quantify brain regions prone to force the transition towards a healthy brain state. Indeed, in the context of whole-brain modelling such external stimulations have been considered^46,47^. Moreover, it has been shown how unclear and often paradoxical the signal that propagates in cortical circuits after non-invasive neurostimulation is^48^. Hence, this further motivates modelling studies in investigating local external stimulations affecting whole-brain changes.

In the Dynamic Sensitivity Analysis paradigm, one important aspect is the description of a brain state. Indeed, how to adequately describe a brain state has been of much focus in modern neuroscience^7^. It is clear that a brain state is linked with the dynamics of the brain, particularly in terms of large-scale network organisations evolving in time, but how to best capture these dynamics across space and time remains a key question^7^. In this study, we chose to summarise brain states in terms of their metastability measure which quantifies the amount of regional variability across the whole-brain and has been shown as a robust differentiator of AD stages^29^. Complimentarily, other measures describing brain states in terms of dynamical system’s theory can be considered, such as the Probabilistic Metastable Substates shown to capture differences between sleep stages^36^ and distinguish responders and non-responders to psilocybin treatment to depression^49^. Alternatively, novel methods from non-equilibrium and turbulent systems have been considered to further expand the notion of a brain state for future in-silico exploration for the Dynamic Sensitivity Analysis^8,50^.

Rebalancing brain states has also received attention from control network theory where strategies are implemented to drive trajectory within a complex system from an initial to a target brain state^51–53^. Such an approach differs from the methods applied in this study where we describe the target state in terms of its dynamical regime with a certain property (here in terms of metastability for example)^54,55^. This is important as in that regard we don’t specify the trajectory but rather approximate the optimal target brain state in terms of its dynamics, and via in-silico perturbation let the system to approximate or rebalance itself towards that dynamic regime.

## Conclusions

In-silico perturbations have the potential to reconceptualise the impacts of stimulation from nodal to whole-brain specific outcomes and guide clinical trials in a more computationally driven way. We used Dynamic Sensitivity Analysis for description, explanation, and prediction to suggest neurostimulation for effective cognitive intervention for Alzheimer’s disease. Here, we demonstrate an individual’s region’s susceptibility to affect global brain dynamics of Alzheimer’s patients towards those observed in healthy controls. This is driven by structural hub regions known to affect the global network in the most impactful way.

## Acknowledgements

Jakub Vohryzek is supported by EU H2020 FET Proactive project Neurotwin grant agreement no. 101017716, Morten L. Kringelbach is supported by the European Research Council Consolidator Grant: CAREGIVING (615539), Pettit Foundation, Carlsberg Foundation and Center for Music in the Brain, funded by the Danish National Research Foundation (DNRF117). Joana Cabral is supported by the Portuguese Foundation for Science and Technology CEECIND/03325/2017, UIDB/50026/2020 and UIDP/50026/2020, Portugal. Gustavo Deco is supported by the Spanish Research Project PSI2016-75688-P (Agencia Estatal de Investigación/Fondo Europeo de Desarrollo Regional, European Union); by the European Union’s Horizon 2020 Research and Innovation Programme under Grant Agreements 720270 (Human Brain Project [HBP] SGA1) and 785907 (HBP SGA2); and by the Catalan Agency for Management of University and Research Grants Programme 2017 SGR 1545.

